# Whole genome sequencing for predicting clarithromycin resistance in Mycobacterium abscessus

**DOI:** 10.1101/251918

**Authors:** Samuel Lipworth, Natasha Hough, Laura Leach, Marcus Morgan, Katie Jeffery, Monique Andersson, Esther Robinson, E. Grace Smith, Derrick Crook, Tim Peto, Timothy Walker

**Author notes:** Contributed equally (Samuel Lipworth, Natasha Hough). Corresponding author, Samuel Lipworth.

## Abstract

*Mycobacterium abscessus* is emerging as an important pathogen in chronic lung diseases with concern regarding patient to patient transmission. The recent introduction of routine whole genome sequencing (WGS) as a replacement for existing reference techniques in England provides an opportunity to characterise the genetic determinants of resistance. We conducted a systematic review to catalogue all known resistance determining mutations. This knowledge was used to construct a predictive algorithm based on mutations in the *erm(41)* and *rrl* genes which was tested on a collection of 203 sequentially acquired clinical isolates for which there was paired genotype/phenotype data. A search for novel resistance determining mutations was conducted using an heuristic algorithm.

The sensitivity of existing knowledge for predicting resistance in clarithromycin was 95% (95% CI 89 – 98%) and the specificity was 66% (95% CI 54 – 76%). Subspecies alone was a poor predictor of resistance to clarithromycin. Eight potential new resistance conferring SNPs were identified. WGS demonstrates probable resistance determining SNPs in regions the NTM-DR line probe cannot detect. These mutations are potentially clinically important as they all occurred in samples predicted to be inducibly resistant, and for which a macrolide would therefore currently be indicated. We were unable to explain all resistance, raising the possibility of the involvement of other as yet unidentified genes.

## Introduction

The *Mycobacterium abscessus complex (M. abscessus)* are rapidly growing nontuberculous mycobacterium (NTM) of increasing clinical concern with a rising burden of associated pulmonary disease (Prevots et al. 2010). *M*. *abscessus* poses a significant problem, particularly in patients with cystic fibrosis (CF), where infection is associated with a more rapid decline in lung function and can be a barrier to transplantation(Esther et al. 2010). Of particular concern are the findings from recent work that have suggested person-to-person transmission of virulent clones amongst the CF population within a healthcare setting (Bryant et al. 2013, 2016), although not all studies have supported this (Harris et al. 2014; Tortoli et al. 2017).

The taxonomy of *M*. *abscessus* is contentious. It is currently divided into three subspecies: *M*. *abscessus* subspecies *abscessus (Mabs)*, *M*. *abscessus* subspecies *massiliense (Mmas)*, and *M*. *abscessus* subspecies *bolletii (Mbol)(Adekambi et al. 2017)*. The organism has intrinsic resistance to multiple antibiotics including β-lactams, rifampicin and aminoglycosides due to the synergistic action of the cell envelope and genetic factors (Nessar et al. 2012). Treatment requires prolonged courses of multiple antibiotics, but outcomes are thought to vary across the different subspecies. *Mmas* has been associated with clarithromycin susceptibility and favourable treatment outcomes, whereas *Mabs* has been associated with inducible macrolide resistance and poorer treatment outcomes (Koh et al. 2011).

Whole genome sequencing has been implemented in stages across England since December 2016, replacing existing reference techniques for mycobacterial identification. As a consequence, there is now the opportunity to explore the molecular determinants of drug resistance for all clinical NTM isolates. Macrolides are important agents in the management of NTM infection, The American Thoracic Society/Infectious Diseases Society of America and British Thoracic Society (ATS/IDSA and BTS) guidelines recommend including a macrolide in treatment regimens where samples are either susceptible, or demonstrate inducible resistance (Haworth et al. 2017; Griffith et al. 2007). They act by binding to the 50S ribosomal subunit and resistance in mycobacteria primarily occurs through target site modification for example by erm methylases and point mutations (Nash, Brown-Elliott, and Wallace 2009).As there is a particularly strong correlation between *in vitro* susceptibility and clinical response to macrolide treatment of *M. abscessus (Jeon et al. 2009; Choi et al. 2017)*, we have undertaken a study to assess the feasibility of predicting clarithromycin susceptibility from whole genome sequencing data for all three subspecies of *M*. *abscessus*.

## Methods

### Literature search

We first conducted a systematic review of the literature to search for known drug resistance conferring mutations in *M*. *abscessus*. Pubmed was searched with the terms‘*Mycobacterium abscessus*’ AND ‘clarithromycin’ OR ‘macrolide’ OR ‘drug resistance’ OR ‘antibiotic resistance’, looking for English language articles published up to April 2018. To be included in the final list, articles had to contain genotyping of coding regions relevant to clarithromycin resistance in *M*. *abscessus* in addition to paired drug susceptibility data. Studies looking at both clinical and non-clinical samples were included. 298 abstracts were screened for relevance and 81 full text articles were obtained of which 26 met the inclusion criteria (Figure 1).

**Figure 1:**
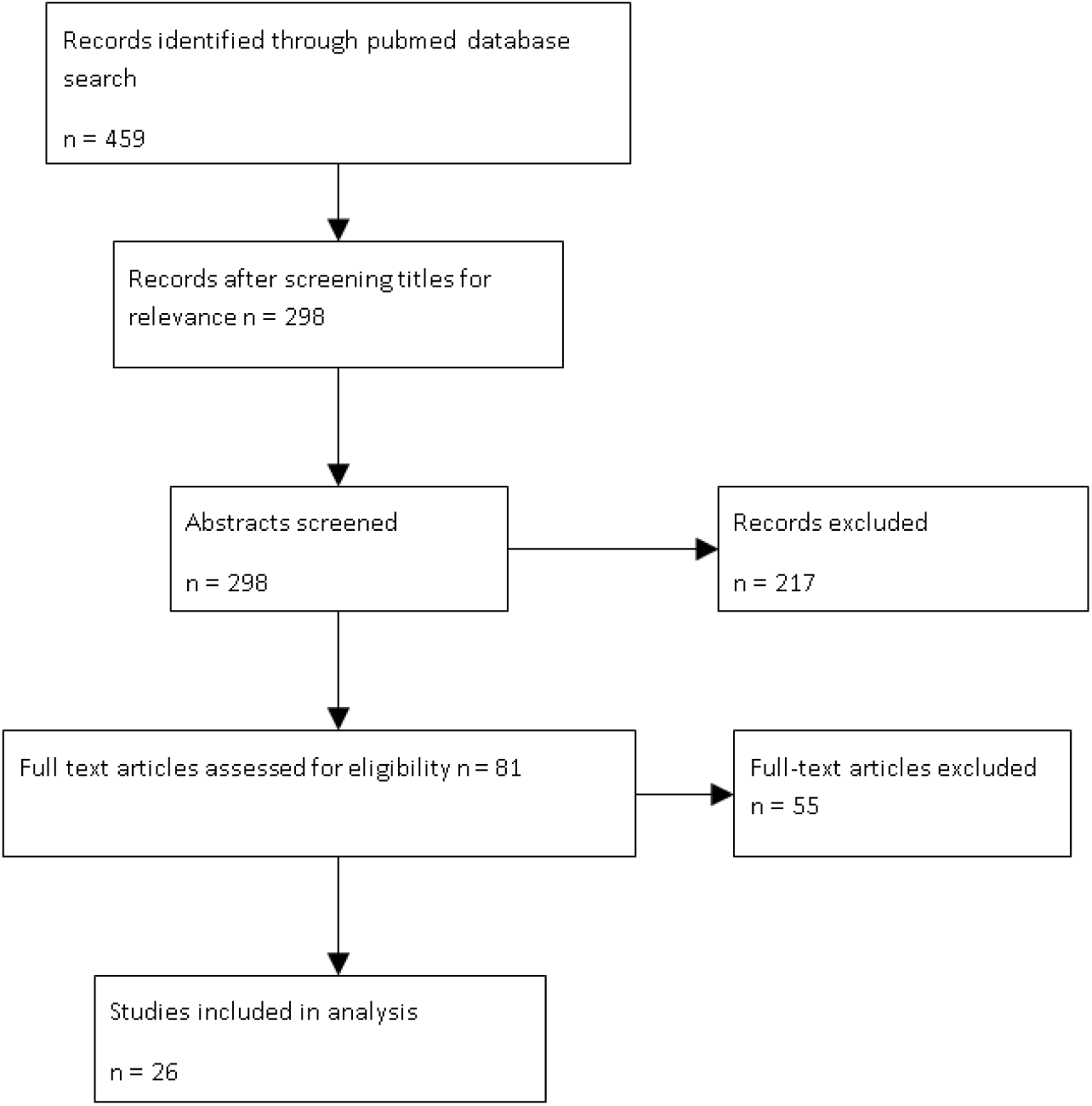
Flow diagram showing stages of the systematic literature search

### Sample selection and sequencing

We next sought all available clinical isolates (N = 180) which had undergone whole genome sequencing by the Public Health England (PHE) laboratory in Birmingham (UK) as part of the routine diagnostic workflow, and for which paired phenotypic data were also available. We supplemented this with 23 isolates for which the same data was available from a WGS archive at the University of Oxford. Isolates were collected between May 2014 and January 2017 and no prior selection according to site of isolation nor whether confirmed M. abscessus complex disease by guidelines was made. Clinical samples were cultured in BD BACTEC^™^ MGIT^™^ liquid mycobacterial growth indicator tubes from which an aliquot was removed to be prepared for WGS as previously described (Votintseva et al. 2015).

Libraries for Illumina Miseq sequencing were prepared using the Illumina Nextera XT protocol with manual library normalisation. Samples where batched 12 to 16 per flow cell and paired end sequencing was performed with the MiSeq reagent kit V2. Bioinformatics was performed using the PHE bioinformatics pipeline as previously described (Walker et al. 2015; Votintseva et al. 2015). Briefly, reads were mapped to the *Mabs* reference genome (NC_010397.1) with Stampy v1.22 and variants called using Samtools v0.1.18 (Only variants with >= 5 high-quality reads, mean quality per base >= 25 and > 90% high-quality bases were retained as variants; heterozygous variants with >10% minor variant were not retained). A self-self blast approach was used to mask repetitive regions. Sub-species were identified by computing maximum likelihood (ML) phylogenetic trees incorporating published representative isolates from each subspecies. A whole genome SNP alignment was used as input to IQ-TREE OMP version 1.5.5 using a generalised time reversible model. The *erm(41)* and *rplV* genes were manually inspected for insertions/deletions from aligned fasta files using Seaview version 4.6.2. All newly sequenced data has been uploaded to NCBI under project accession number PRJNA420644.

### Drug susceptibility testing

Phenotypic drug susceptibility testing (DST) was performed at the PHE National Mycobacterial reference service in London. DST was performed using the broth microdilution method with 96-well RAPMYCO microtitre plates (Mueller Hinton medium with TES buffer, Thermo Fisher). Plates were read at day three post-inoculation, and if poor growth again at day 5, according to Clinical and Laboratory Standards Institute (CLSI) guidelines (Committee for Clinical Laboratory Standards 2000). Isolates deemed susceptible or intermediate were re-incubated and read again at days 7, 14 and 21. Those found to be resistant (R MIC □ 8 μg/ml) at any of these time points are described as phenotypically resistant. A call of phenotypically sensitive (S MIC □ 2 μg/ml) or intermediate (I MIC > 2 - < 8 μg/ml) was only made after the full 21 days of incubation. This study was an opportunistic retrospective analysis of routinely collected clinical data and as such phenotypic testing was not repeated on discordant isolates.

### Genotypic prediction of clarithromycin susceptibility

We used BioPython software to extract base calls from whole genome sequence FASTA files, comparing these to a list of genomic loci which our literature search indicated were associated with clarithromycin resistance (table 2). We then predicted phenotypes using an hierarchical algorithm (Figure 2). A resistant phenotype was predicted where any mutations were present at *rrl* positions 2270 or 2271 (*E*. *coli* numbering 2058/2059), or where the less well characterized *rrl_A2269G or rrl_A2293C or rrl*_G2281A mutations were seen. In the absence of these mutations, susceptibility was predicted where an isolate had a truncated *erm(41)* gene or a C nucleotide at position 28 in *erm(41)*. Inducible resistance was predicted where a wild type call (T) was present at position 28 in *erm41*. However, if an *erm41*_C19T mutation was also present, susceptibility was predicted instead of inducible resistance. In cases where there was a null call at *rrl* 2270/2271, we subsequently attempted local assembly of the *rrl* gene using Ariba (Hunt et al. 2017), followed by comparison by alignment against the reference. Where this was not possible due to low coverage in this region, no prediction was made. Statistics quoted were calculated using R Studio v1.1.383.

**Table 1.** -summary of genotypes and corresponding clarithromycin phenotypes for the 203 isolates. Pos - *M*. *abscessus* numbering position in gene, prediction - genotypic prediction using the algorithm shown in Figure 2, Mabs - *M abscessus absecssus*, Mbol - *M. abscessus bolletii*, Mmas - *M. abscessus massiliense*, N – total number of isolates with genotype, S – Sensitive, R – Resistant, IR – Inducible Resistance

|  | erm41 pos. 28 | erm41 length | erm41 pos. 19 | rrl. pos 2269 | rrl pos. 2270 | rrl pos. 2271 | N | Phenotype | Prediction |
| --- | --- | --- | --- | --- | --- | --- | --- | --- | --- |
| **Mabs** | T | full | C | A | C | A | 5 | 5R | R |
|  | T | full | C | A | T | A | 2 | 2R | R |
|  | C | full | C | A | A | A | 37 | 4I, 2R, 31S | S |
|  | T | full | C | A | A | A | 87 | 62R, 25S | R |
|  | Excluded due to inadequate coverage over rrl 2270-2271 |  |  |  |  |  | 12 | 10R 2S |  |
| **Mbol** | T | full | C | G | A | A | 1 | 1R | R |
|  | T | full | C | A | G | A | 2 | 2R | R |
|  | T | full | C | A | A | A | 13 | 11R, 2S | R |
|  | Excluded due to inadequate coverage over rrl 2270-2271 |  |  |  |  |  | 4 | 4R |  |
| **Mmas** | T | truncated | C | A | C | A | 3 | 3R | R |
|  | T | truncated | C | A | G | A | 3 | 3R | R |
|  | T | truncated | C | A | A | G | 5 | 5R | R |
|  | T | truncated | C | A | A | A | 24 | 21S, 3R | S |
|  | T | full | C | A | A | A | 1 | 1R | R |
|  | Excluded due to inadequate coverage over rrl 2270-2271 |  |  |  |  |  | 4 | 4R |  |

**Table 2:**
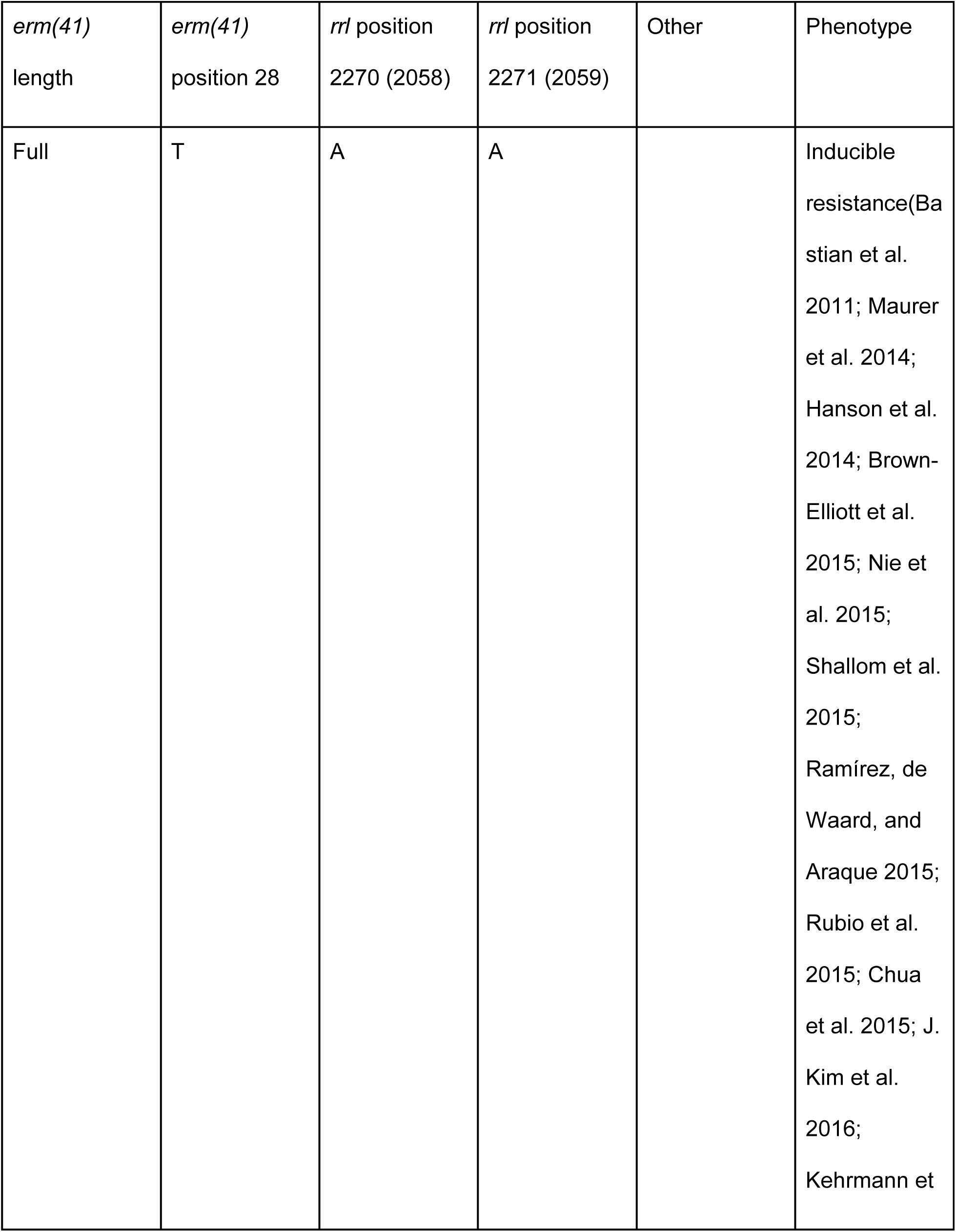
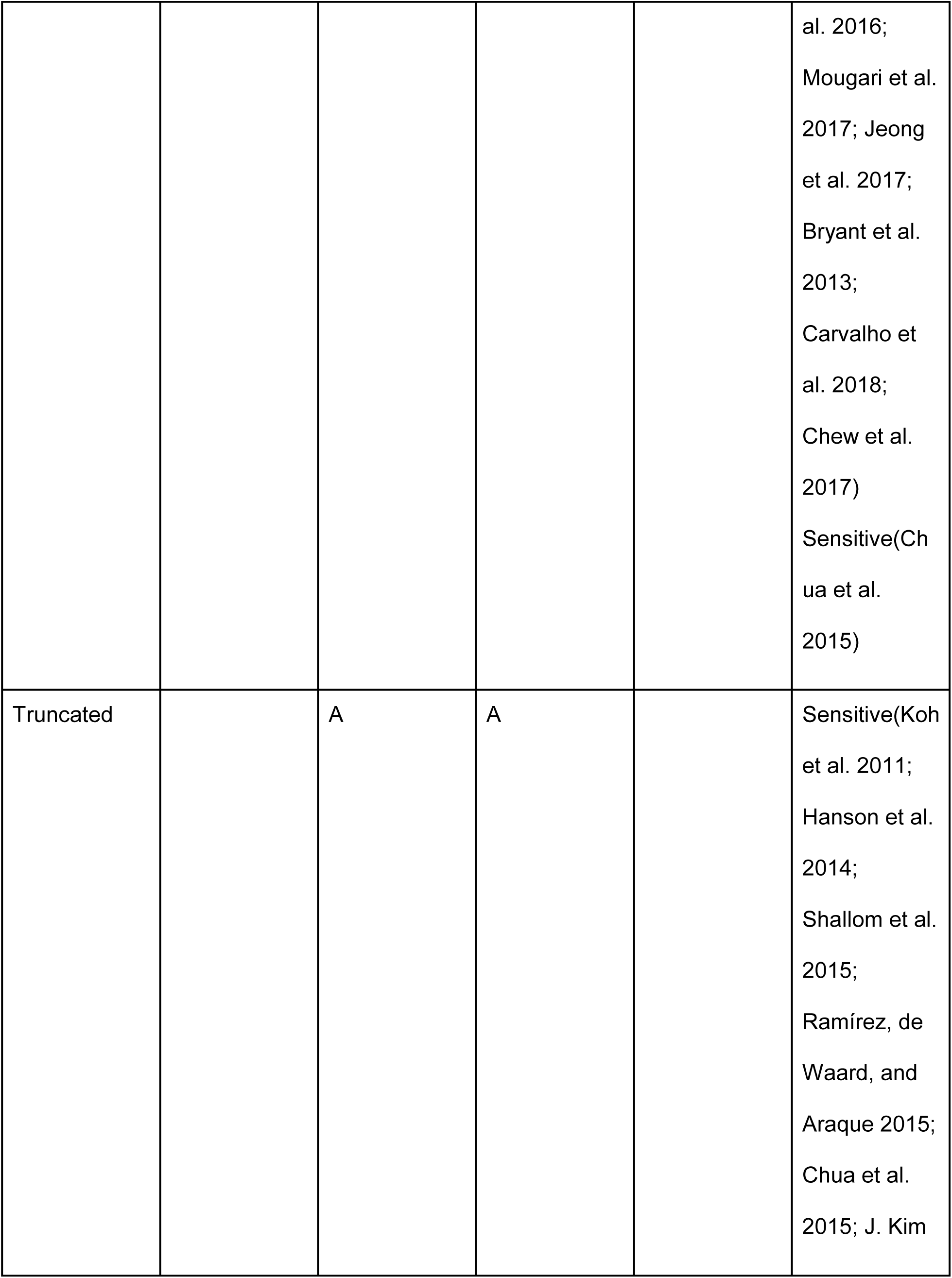
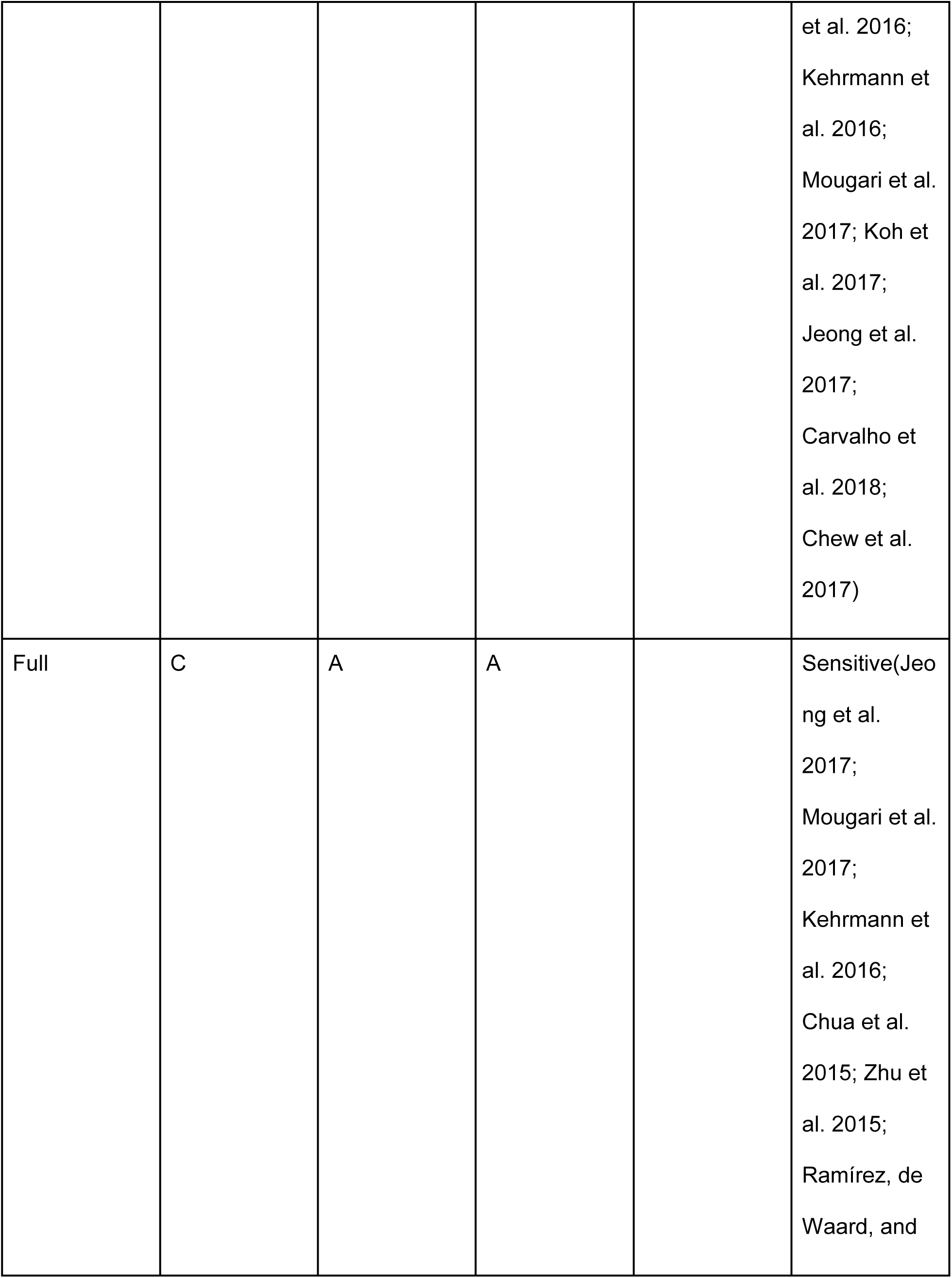
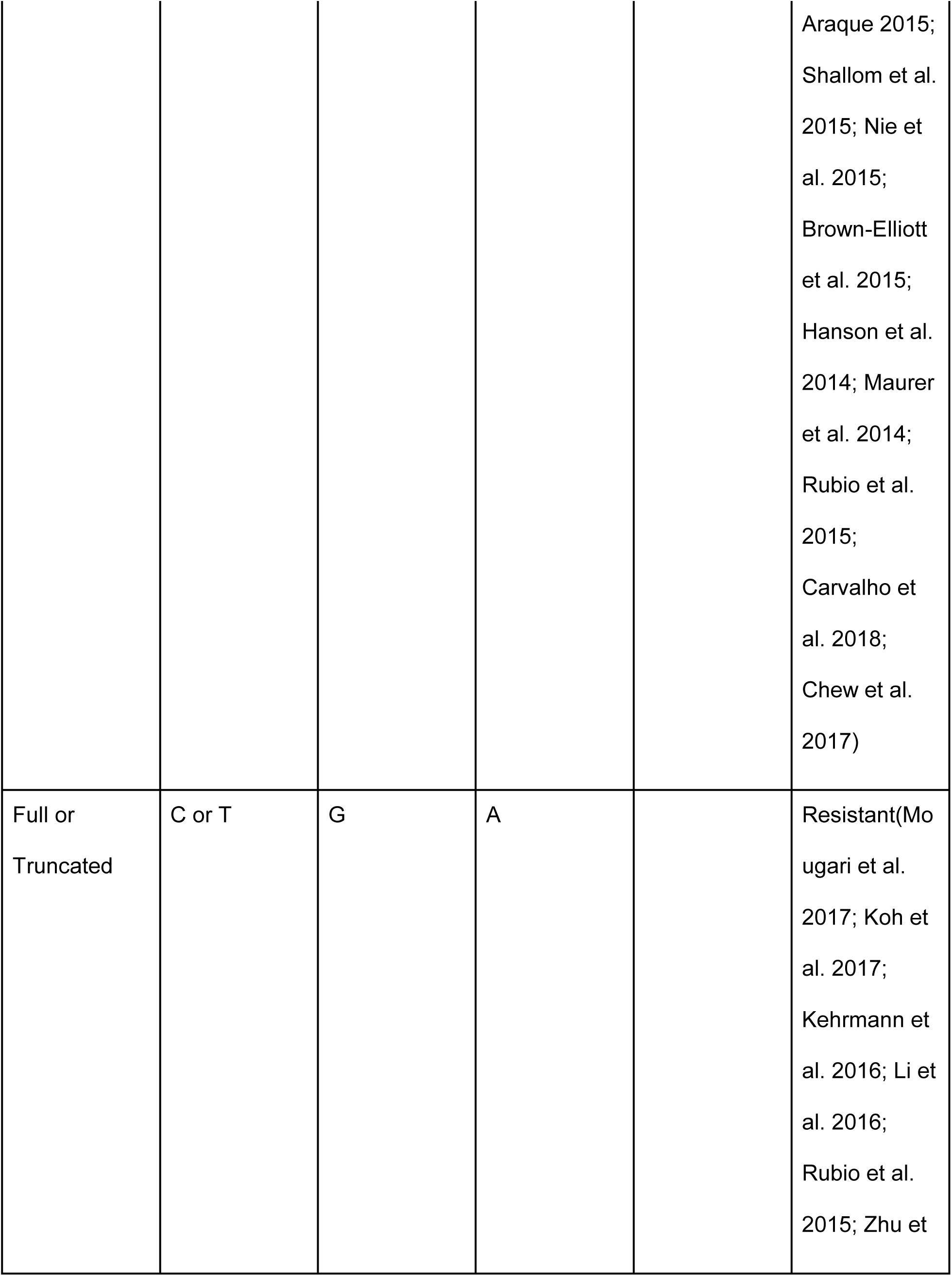
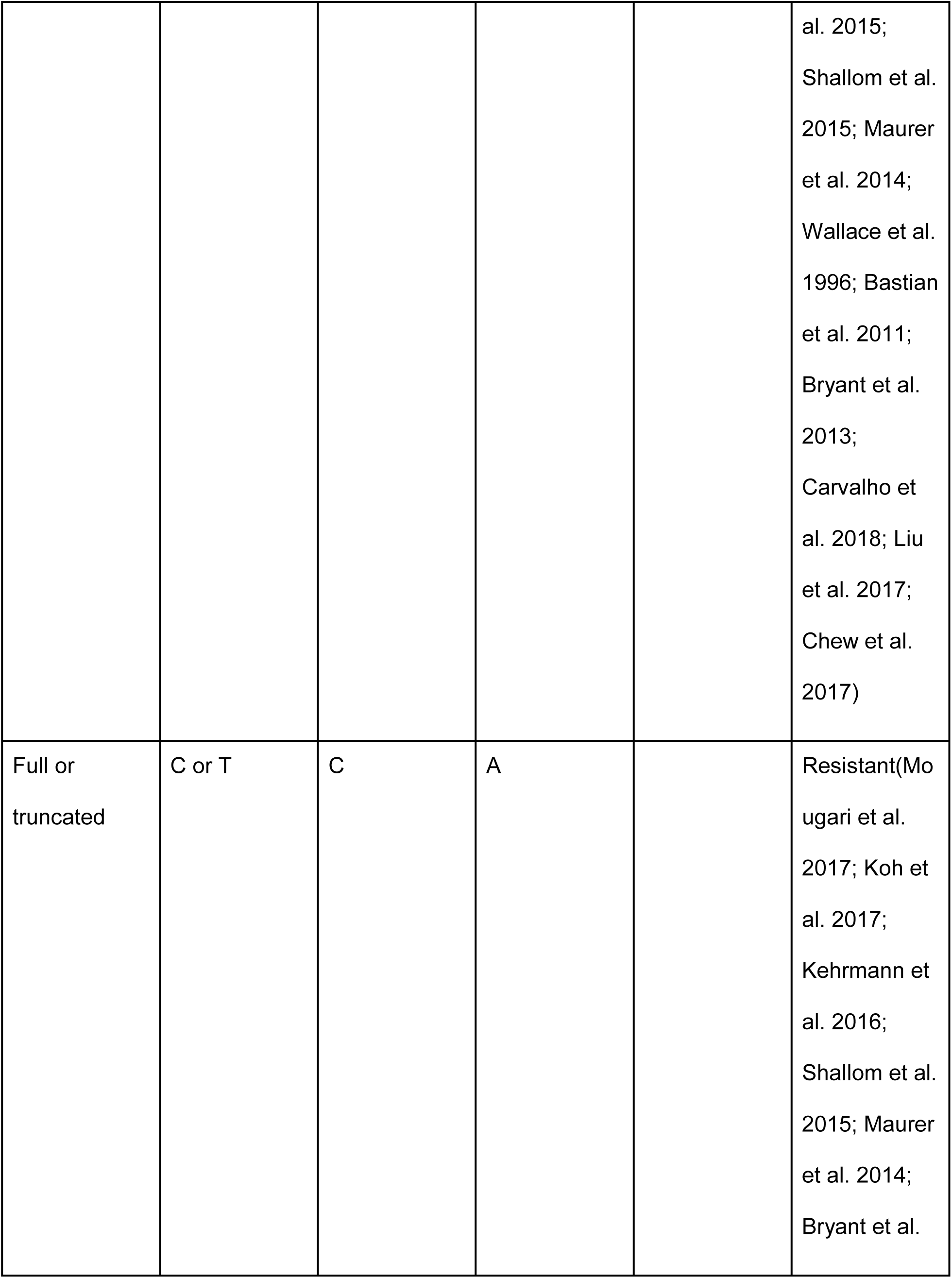
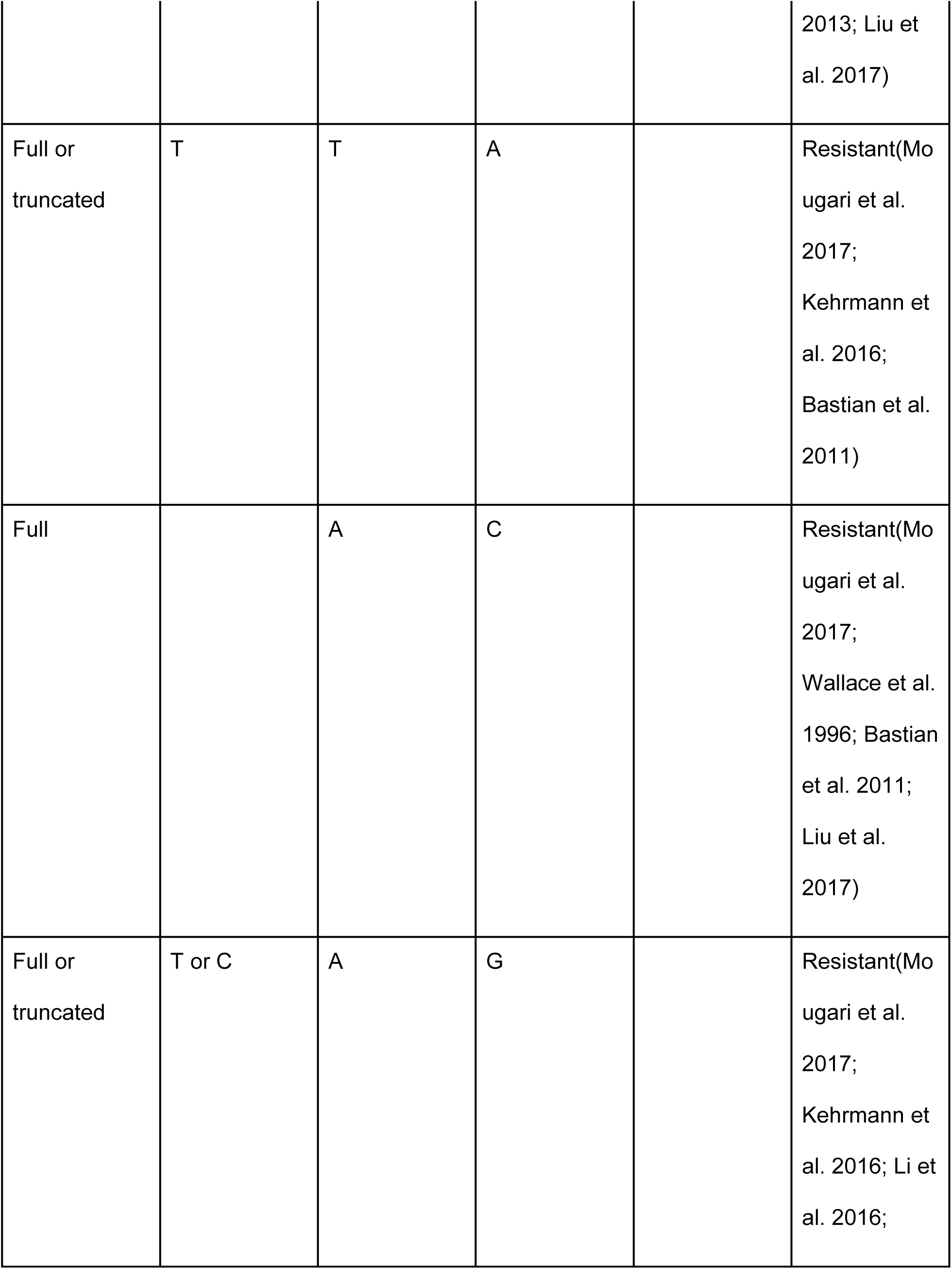
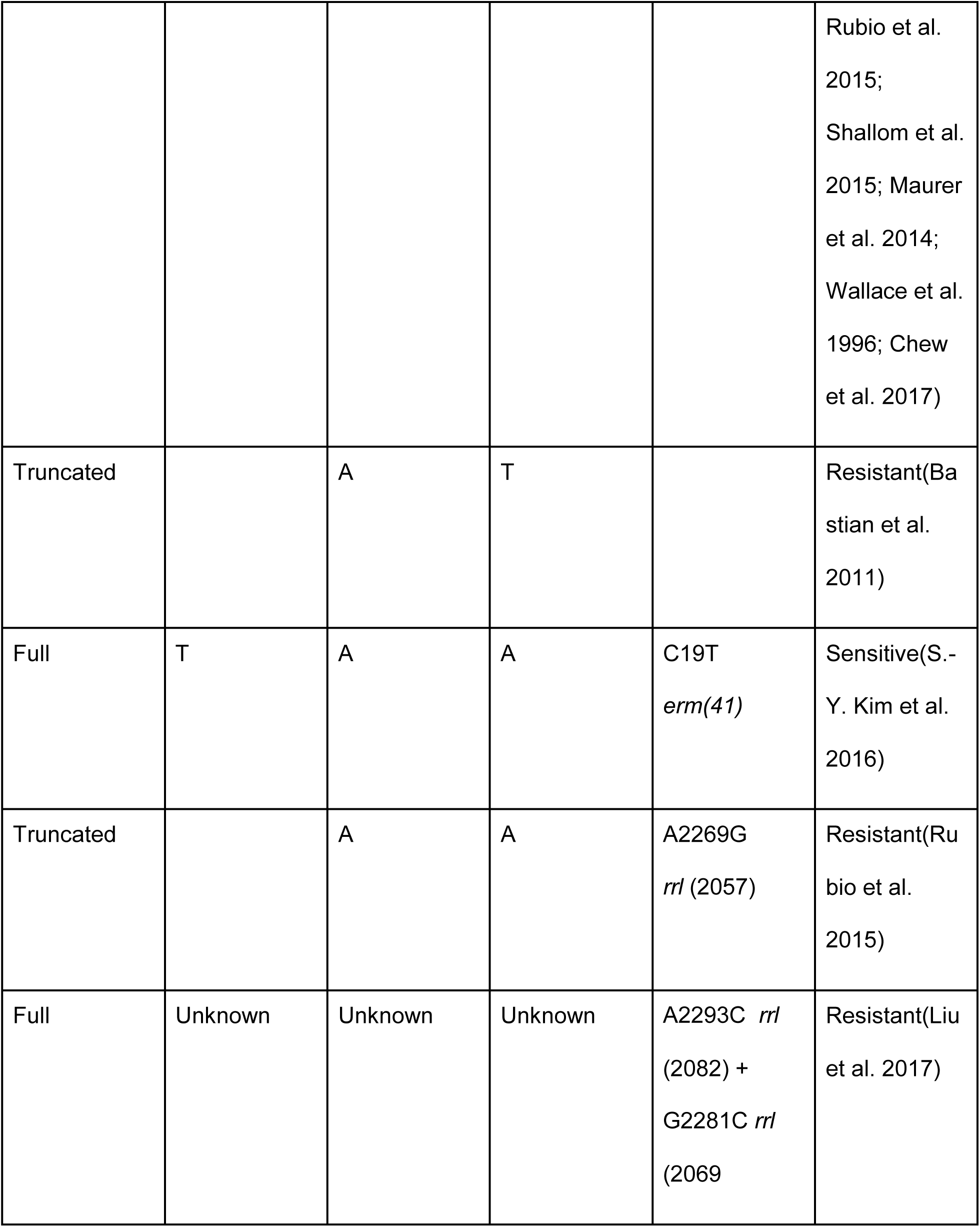
Resistance determining mutations for clarithromycin identified in the literature search. *M*. *abscessus* numbering is used with *E. coli* numbering in brackets.

**Figure 2:**
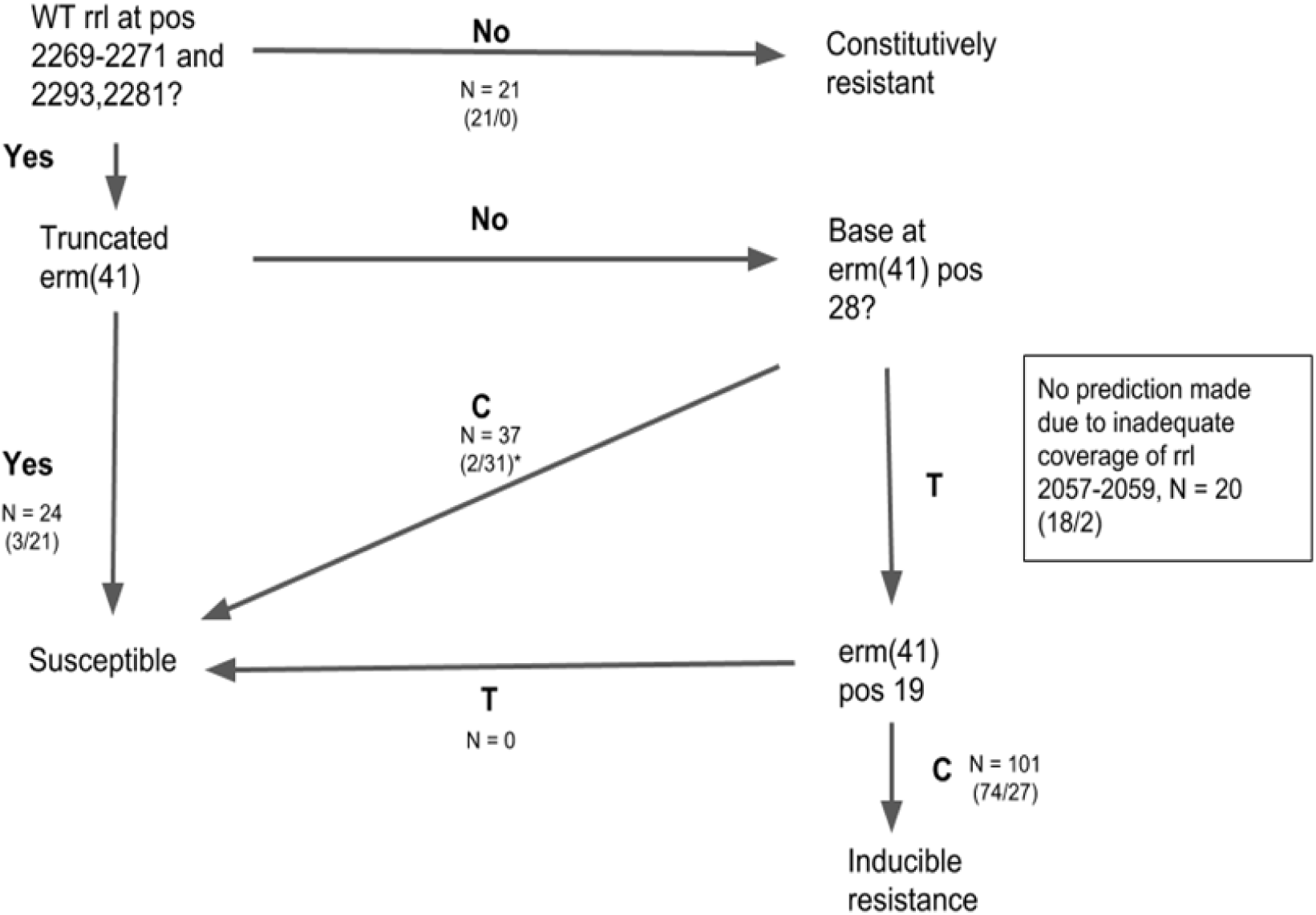
decision algorithm for predicting drug resistance in *M*. *abscessus* based on the literature search with numbers of isolates meeting each predictive criterion shown. Bracketed numbers represent (N resistant / N sensitive). ^∗^ 4 isolates had intermediate susceptibility

### Search for novel resistance conferring mutations

We attempted to characterise new resistance mutations within genes linked to drug resistance from the literature search. To maximise power for discovering new potential resistance mutations, we included all genomes available to us. All variants in these genes or their promoter regions were extracted using Python software from variant call files. Phylogenetic SNPs were identified by considering each subspecies in turn, assumed to be benign and excluded from further analysis.

We considered variants at the level of SNPs in promoter regions or rRNA and amino acid changes in coding regions. A mutation (a variant with an observable phenotype) was characterised as causing resistance if it occurred as the only variant in the relevant region in a resistant isolate or if it was always associated with resistance when it was seen and did not co-occur with any other mutations known to cause resistance. Variants were characterised as consistent with susceptibility (‘benign’) if all isolates were susceptible when it occurred alone or if it occurred only in susceptible isolates. We assumed no prior knowledge in this section of the analysis and the identification of known resistance SNPs was used as an internal validation of our approach.

## Results

We studied 143 *Mabs*, 20 *Mbol* and 40 *Mmas* genomes. Genotypic predictions were made on the basis of mutations identified by the literature search. All relevant mutations identified were contained in the genes *rrl* and *erm(41)* (figure 2 and table 2). The genes *rplV*, *whiB7* and *rpld* were also considered of potential interest and were additionally searched for variants.

### Genotypic predictions

Inducible resistance was predicted in 101 isolates, of which, 74/101 (73%) were reported as phenotypically resistant. After excluding isolates for which no prediction could be made due to missing data in key genomic loci (n = 20) as well as those with an intermediate phenotype (n = 4), the sensitivity was 95/100 (95%, 95% CI 89 - 98%) and specificity was 52/79 (66%, 95% CI 54 - 76%) . The very major error rate (phenotype resistant, WGS prediction sensitive) was 5/100 5% (95% CI% 1 – 9%) and the major error rate (phenotype susceptible, WGS prediction resistant) was 27/79, 34% (95% CI 24 – 44%). Positive predictive value was 95/122, 78% (95% CI 69 - 85%) and the negative predictive value was 52/57, 91% (95% CI 81 - 97%) (Table 3). The F score for WGS predictions was 0.86. When isolates with a prediction of inducible resistance were further excluded, the specificity of a resistance prediction was 21/21 (100%, 95% CI 93 - 100%) and the sensitivity was 21/26 (81%, 95% CI 61 - 93%).

**Table 3.** -WGS Predictions vs DST phenotype for clarithromycin. Sensitivity/Specificity/PPV/NPV are calculated excluding isolates with and intermediate phenotype and those where no prediction was made due to inadequate coverage at key positions.

|  | In vitro phenotype |  |  |
| --- | --- | --- | --- |
| Genomic Prediction | Sensitive | Resistant | Intermediate |
| No Prediction* | 2 | 18 | 0 |
| Inducible resistance | 27 | 74 | 0 |
| Resistant | 0 | 21 | 0 |
| Sensitive | 52 | 5 | 4 |
| Sensitivity | 95% (95% CI 89 - 98%) |  |  |
| Specificity | 66% (95% CI 54 - 76)% |  |  |
| Positive predictive value | 78% (95% CI 69 - 85%) |  |  |
| Negative predictive value | 91% (95% CI 81.0 - 97%) |  |  |

### Clarithromycin resistance in the subspecies

81/143 *Mabs* were resistant, 58 sensitive and 4 intermediate. For *Mbol* 18/20 were resistant and for *Mmas* 19/40 were resistant (table 1). There was one *Mmas* isolate carrying a full length *erm(41)* gene which was phenotypically resistant to clarithromycin. This was not unexpected from a genotypic perspective as it harboured a wild type thymine nucleotide at position 28 *erm(41)*, associated with inducible resistance.

### Mechanisms of resistance

The negative predictive value of a truncated *erm(41)* gene for clarithromycin susceptibility was 53% (21/39 - there was one *Mmas* isolate with a full length *erm(41))*. In 11/18 instances, resistance in the presence of a truncated *erm(41)* could be explained by a mutation in position 2270 or 2271 in *rrl*. No coverage at all was seen at these positions for 4/18 isolates. No genomic explanation could be identified for the remaining three discordant isolates (table 1).

All isolates which had any mutation of positions 2269, 2270 or 2271(E. Coli numbering 2057, 2058, 2059) in *rrl* were resistant to clarithromycin (21/203 (10%)). Such a mutation was found in 3 *Mbol*, 11 *Mmas* and 7 *Mabs* isolates. We did not observe any isolates with an *rrl* mutation which also harboured a T28C mutation in *erm(41)*. Where this occurred in isolates reported in the literature, they were always resistant (Kehrmann et al. 2016; Rubio et al. 2015).

Of 37 isolates with a T28C mutation in *erm(41)* and no other relevant mutations, 84% (31/37) were susceptible to clarithromycin, 11% (4/31) had intermediate susceptibility and 5% (2/31) were resistant. This mutation was exclusively found in *Mabs* isolates. We did not identify any drug resistance associated mutations in any of these intermediate or resistant isolates. Across all three subspecies, of 101 isolates with the T28_*erm41* call associated with inducible resistance (and no other relevant mutation), 73% (74/101) were resistant and 27% (27/101) susceptible at the final day 21 reading.

### De novo search for resistance determining mutations

The search for potential novel resistance determining mutations for clarithromycin revealed 13 SNPs of interest (table 4). Of these, five have previously been described in the literature. There were additionally four SNPs (*rrl*_A2746T, *rrl*_G836A, *rrl*_T2674G and *rrl*_T636C) which were only ever seen in resistant isolates but always co-occurred with known resistance determining SNPs. There was one phenotypically resistant isolate which harboured 18 novel SNPs. On performing a nucleotide BLAST of a 120 base region encompassing all of these SNPs, there was a 99% (E 2 ×10^-53^) match with *Streptococcus species*. This therefore likely represents sample contamination with flora from the nasopharynx. No new resistance associated variants were discovered in *rplV*, *rpld* or *whiB7*.

**Table 4:** Mutations (both novel and previously described) detected during *de novo* search for resistance determining SNPs. Rule 1 = occurs as only SNP in relevant regions in resistant isolate, rule 2 = all samples resistant when SNP occurs, never seen in sensitive isolate. All numbering is relative to M. abscessus. ^∗^ mutation already described in literature - *M. abscessus rrl* numbering 2270/2271 is *E. coli* numbering 2058/2059. ^∗∗^ mutation in *erm(41)* promoter region, 31 bases upstream of start of coding region.

| Position | Nucleotide/Amino acid change | Rule met |
| --- | --- | --- |
| rrl 2039 | A > G | 1 |
| rrl 1401 | T > C | 2 |
| rrl 371 | T > C | 2 |
| rrl 795 | G > A | 1 |
| rrl 2270* | A > C | 1 |
| rrl 2270* | A > G | 2 |
| rrl 2271* | A > G | 2 |
| rrl 2270 * | A > T | 2 |
| erm(41) 131 | A > V | 2 |
| rrl 2279 | G > A | 2 |
| rrl 2269* | A > G | 2 |
| erm (41) -31** | A > T | 2 |
| rrl 1932 | A > G | 2 |

## Discussion

We conducted a systematic review of drug resistance determining mutations for clarithromycin in *M*. *abscessus* and used the results to make genotypic predictions. The sensitivity of this approach was 95% (95% CI 89 – 98%) and the positive predictive value 78% (95% CI 69 – 85%). The prevalence of resistance amongst our collection of isolates was high compared to that which has been reported elsewhere (Koh et al. 2011; Hatakeyama et al. 2017; Li et al. 2016; Cowman et al. 2016).

These results show that for clarithromycin, drug resistance can be predicted from WGS data as it has been previously through targeted PCR and line probe assays such as the Hain GenoType NTM-DR. Assessment of the genotype of *erm(41)* with molecular diagnostics allows prediction of its functional status which has been thought to correlate to treatment outcome (Haworth et al. 2017). Similarly, as the absence of a functional *erm(41)* gene has been associated with good therapeutic outcomes its molecular detection ought to be beneficial to patients (Koh et al. 2011), although in our study this alone was not an adequate predictor of in vitro resistance. A genotypic prediction of inducible resistance produced a variable phenotype in our study (27/101 sensitive). Discriminating such isolates predicted to be inducibly resistant which are unexpectedly sensitive after prolonged incubation with clarithromycin or show early time point high level resistance may help to identify additional genotypic markers to better identify patients more likely to benefit from the use of macrolides…

In addition to mutations identified in the literature search, we also managed to identify variants that may plausibly be new resistance determining mutations. However, these will require validation against an independent data set. Using routinely collected diagnostic data to improve our understanding of the molecular determinants of drug resistance is a key advantage WGS has over line-probe assays or PCR. The eight previously undescribed mutations we report in this work could be of clinical importance because they all occur in samples which the existing literature predicts to be inducibly resistant. As BTS guidelines recommend that patients with such isolates should be given a macrolide, it is important to determine further whether these SNPs are true resistance-determinants, and whether macrolide therapy should be avoided in their presence.

Previous authors have suggested that it is clinically useful to discriminate between subspecies, (Koh et al. 2011) as *Mmas* is typically associated with durable susceptibility to clarithromycin and *Mbol* and *Mabs* with inducible resistance (unless the T28C mutation is present). We found identifying sub-species alone to be an inadequate predictor of *in vitro* clarithromycin phenotype. There were three *Mmas* isolates in our collection that were resistant to clarithromycin and had no mutations known to be relevant. Mougari and colleagues found that in 39/40 *Mmas* selected for clarithromycin resistance, this could be explained by an rrl mutation at positions 2270/2271 with a further sample containing an *rplV* insertion (Mougari et al. 2017). All of our isolates contained this ‘insertion’ (also present in the NC_010397.1 reference) which was associated with susceptibility to clarithromycin except in the presence of a relevant rrl mutation.

In keeping with previous reports, we identified an isolate of *Mmas* with a full length *erm(41)* and a thymine nucleotide at position 28(Shallom et al. 2015). This likely represents recombination between the subspecies. A recent study showed the Hain GenoType NTM-DR line probe assay incorrectly predicted subspecies in 8% of samples, presumably because it lacks the whole genome resolution provided by sequencing and is vulnerable to between species recombination (Kehrmann et al. 2016).

Despite analysing all mutations occurring in *erm(41)* and *rrl* for the full collection of genomes, we were unable to predict all clarithromycin resistance. This may be because there are other genes implicated or due to unreliable DST results. Future work should aim to select discordant genotypes and identify additional infrequently occurring genetic loci implicated in clarithromycin resistance, for example by using genome wide association (GWAS) approaches. All of the new clarithromycin resistance mutations we discovered occurred in isolates which we originally predicted to be inducibly resistant. Although M. abscessus is primarily thought to be an environmental organism, these patients may be colonised for long periods with subsequent potential exposure to multiple courses of macrolides. An alternative hypothesis may therefore be that some or all of these SNPs are compensatory mutations which act to reduce a fitness cost of the expression of *erm*, which has been experimentally demonstrated in other bacteria (Gupta et al. 2013). There were four SNPs which only occurred in resistant samples but were always seen with a known drug resistance causing SNP, possibly also representing compensatory mutations.

Key weaknesses of our study include that we were unable establish a temporal relationship between antibiotic prescribing and inducible phenotypic resistance as we did not have the relevant ethics approval to link to patient records. If for example, any SNPs on our list of novel mutations were observed in isolates from patients who had never previously had macrolide therapy, it would be much more likely that they were genuine resistance conferring rather than compensatory mutations. In addition it is possible that some of the genomes were same patient replicates over a number of months/years, although this may have also diversified the range of mutations observed. We chose to include all available samples to maximise detection of low frequency resistance determining SNPs meaning there was no validation set available. Our list of novel resistance determining SNPs will therefore require validation on an independent dataset before being applied to the clinical setting. We chose to target a select list of genes with known SNPs identified in the literature search; other approaches such as GWAS will likely be additive to the knowledge base we present here.

In summary, WGS allows identification of known resistance conferring mutations as well as demonstrating probable novel resistance determining SNPs in regions the Hain NTM-DR line probe cannot detect which if further validated may change management. Identification of subspecies alone inadequately predicts macrolide resistance in *M. abscessus*. Our data does not support the replacement of phenotypic tests at this point in time; as more paired genome/DST data becomes available in the near future, and we learn more about the molecular determinants of drug resistance, it is likely that sensitivity and specificity of WGS resistance prediction will improve. Given that WGS data is already being produced in the UK for the purposes of molecular epidemiology, it would now be possible to phase out existing molecular tests and replicate their results *in silico* at no additional cost.

### Transparency declaration

The authors have no conflicts of interest to declare.

